# Geospatial statistics of high field functional MRI reveals topographical clustering for binocular stereo depth in early visual cortex

**DOI:** 10.1101/160788

**Authors:** A.J. Parker, G. S. L. Coullon, R. M. Sanchez-Panchuelo, S.T. Francis, S. Clare, D.A. Kay, E. P. Duff, L. Minini, S. Jbabdi, D. Schluppeck, H. Bridge

## Abstract

**Significance statement:** Functional topography is present throughout the cerebral cortex, often in the form of columns or clusters of neurons with similar functional properties within identified cortical regions. Most of the evidence for these structures comes from work with non-human species. Using high-field strength magnetic resonance imaging in living human cortex, the search for this structure is frustrated by the presence of a background of local spatial correlations in the signal across the cortical surface. This structured background, which we term ‘neural dust’, is a form of noise that imposes a fundamental limit on the detection of clustering and topography. We apply a novel analysis approach that quantifies the background of spatial correlations, so that we achieve reliable identification of high-signal clusters of activation in the human cortex at a scale previously only visible with invasive optical imaging in animals. We searched specifically in visual cortex for correlates of binocular stereoscopic depth. We show both that signals for stereoscopic depth are clustered into specific zones within the cortex and that these signals occur within spatially extended cortical domains, which have similar preferences for the stereoscopic depth. The revealed structures are reliably identified in all subjects tested and in repeated testing of individual subjects. Our methods provide an objective approach for defining the size and locations of clusters of activation within functional images of neural activity.

**Abstract:** A characteristic principle of the organization of cerebral neocortex is the presence of clusters or columns of neurons with similar functional properties. Animals with forward-facing eyes exploit slight differences between the images in the two eyes to determine binocular depth using stereopsis. Evidence for an organized structure in human visual cortex for the representation of stereoscopic depth has proved elusive. Using 7-tesla functional MRI, with gradient-echo echo-planar imaging at 0.75 mm isotropic resolution and a novel analytical approach based on geospatial mapping methods, we find that clustered responses for disparity-defined depth can be clearly segregated from a background of spatially correlated signals in all subjects tested. High-signal clusters are associated with cortical domains as large as 12-15mm across the cortical surface, in which nearby points in the cortical map tend to respond to the same stereoscopic depth. These domains are found predominantly within visual cortical area V3A.

## Introduction

Neuroscientific and clinical investigations over several centuries have revealed the functional organization of the brain at the level of individual cortical areas^1^. However, to determine the mechanisms underlying cortical function, it is necessary to consider the much finer scale of organization that occurs within single cortical areas. At this scale, the functional properties of neurons are often organized systematically with respect to the neurons’ position within the cortical sheet of grey matter. For example, the topographic organization of orientation selectivity of single neurons in primary visual cortex of cat and monkey has a columnar organization: the angular preference of neurons for the orientation of visual bar stimuli changes systematically with position across the cortical sheet, but when moving from the surface to white matter normal to the cortical sheet, the neurons have similar orientation preferences^2-4^. This truly columnar organization may be contrasted with a local clustering of neural properties, in which the neural responses from a particular cortical location tend to have similar characteristics but there are no rules or principles governing the relationships of one cluster to another.

The investigation of within-area organization, particularly in the visual system, has become an important goal in non-invasive human imaging, to match what has been achieved by animal studies. The capacity of functional magnetic resonance imaging (fMRI) to reveal structures that are known to be on the scale of a few mm is limited by a number of factors, even when the greater resolution of high field strengths can be employed^5^. To make deductions about the underlying neuronal structure, it is also necessary to disentangle different elements of the MRI signal, so that neural events in the signal can be isolated from the properties of the blood-oxygen level dependent (BOLD) response. The large gain in BOLD contrast-to-noise ratio (CNR) at 7T allows a significant decrease in voxel volume and thus a consequent increase in spatial resolution relative to 3T. However, inspection of the resulting imaging data shows that even at baseline, and despite the increased sampling resolution, there are substantial spatial correlations in the BOLD signal^6-9^, meaning that a white-noise model is inappropriate. These spatial correlations inevitably limit the ability of high-field imaging to resolve small patches of cortical activation^8^.

The spatial correlations in the background signal also generate apparent structure, which confuses the search for true clustering or topography, particularly when the spatial scale of the correlations is similar to that of the expected size of column-like structures. Although this spatially structured background is a form of noise, it probably arises from a number of weak signals similar to the stronger signals that we wish to identify. For this reason, we refer to this structured background as ‘neural dust’, i.e. something that needs to be cleaned away before we can properly see the neural signal.

Stereoscopic depth perception arises from the slightly different viewpoints of the two eyes, which results in small differences (disparity) in the locations of a visual feature within the retinal images of the left and right eyes. Neurons selective for binocular disparity are found throughout the non-human primate visual system^10^. However, neurons in the primary visual cortex seem to reflect initial stages of processing disparity rather than stereoscopic perception^10-13^. Consistent with this evidence, V1 shows, in many species, a functional organization for eye of origin (ocular dominance columns) and orientation, but there is no known pattern for binocular disparity. By comparison, in extrastriate cortex of macaque, optical imaging reveals a near-to-far organization for disparity within the thick stripes of V2^14^ and neurophysiology of extrastriate areas V2^15^, V3/V3A^16^ and V5/MT^17^ show evidence of clustering of neurons with similar disparity preferences.

The use of functional magnetic resonance imaging (fMRI) in the human visual cortex necessarily yields macroscopic measurements of cortical responses to binocular disparity. In previous studies, this often involved the summation of responses from all voxels falling within entire visual areas, each defined independently by retinotopic mapping. This approach has identified preferential activation by binocular disparity in dorsal occipital regions, such as V3A and the kinetic-occipital area (KO), in addition to multiple subdivisions of the intraparietal sulcus (IPS)^18-23^. Over the past 5-10 years, ultra-high field fMRI has been used to demonstrate both ocular dominance columns and orientation pinwheels in human V1^24,25^. There is also evidence for clustering of motion responses in V5/MT using high resolution imaging at 7T^26^.

Over the past two years, two studies have investigated the fine-scale organization of disparity, one focusing on V3A and KO^27^, and the other on V2 and V3^28^, both providing some evidence for clustering of disparity responses. To search for clustering of disparity tuning in early visual cortex (V1, V2, V3 and V3A), we employed high spatial resolution fMRI of the human occipital lobe (0.75 mm isotropic resolution) at 7T. We used a phase-encoded paradigm in which disparity was changed steadily over time in each stimulus cycle. The response patterns showed significant clustering and topographical structure, which were tested against a correlated noise baseline using novel analytical methods derived from geographical and environmental statistics. We show how these methods provide a rigorous basis for detecting the presence of spatial structure in experimental neuroimaging data.

## Results

Initially we validated our stimulus display and high-field imaging system by replicating an earlier study^21^, in which the specialization of different visual cortical areas for stereoscopic disparity was probed with a stimulus displaying a tiled set of planes at different depths (Fig 1A). The binocular disparity of the planes changed each second during the 16 second viewing period. BOLD imaging responses were collected at 7T field strength using 3D echo-planar scan sequence with 0.75mm isotropic voxel resolution and an overall scan time of 4 sec (see Methods).

**Figure 1:**
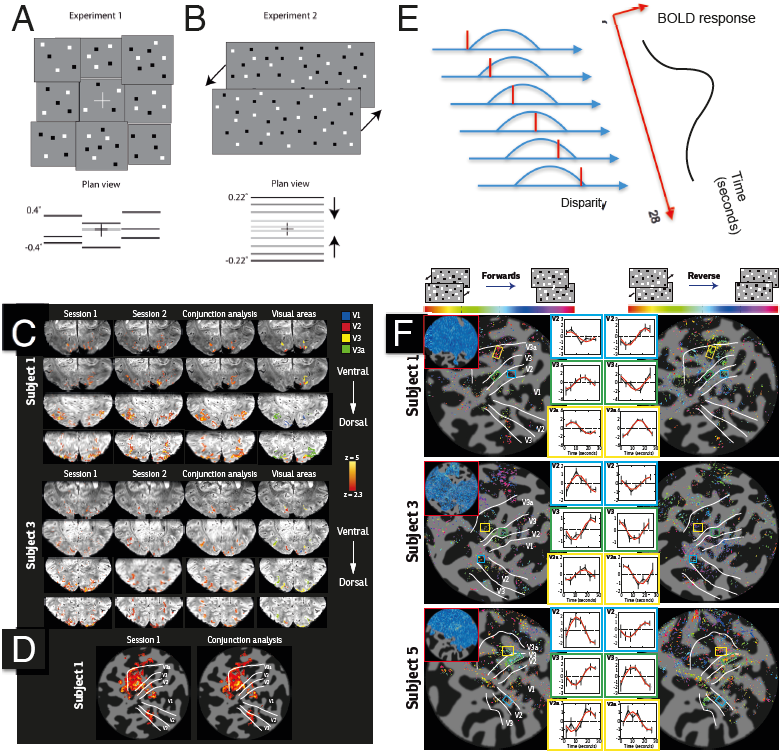
Stimulation of visual cortex with binocular depth patterns leads to specific clusters of responding elements using fMRI at 7 tesla. A: random-dot tiles of different binocular depths alternated with a binocularly uncorrelated field of dots. B: binocular random dot patterns define planes of dots slowly moving in depth from near to far; the upper and lower portions of the pattern move with a sawtooth waveform in opposite directions to ensure there was no net drive to convergence eye movements. C: BOLD fMRI activations of occipital cortical slices measured at 7T field strength in response to pattern A (disparity vs uncorrelated). D: BOLD fMRI responses rendered on computationally flattened cortical surface with single scan (left) and conjunction of two repeated scans (right); there is a strong concordance. E: right panel shows schematic of predicted cortical activations in response to stimulus B, while left panel shows the progression of the disparity plane (red line) over time against the presumed response profile of a cluster of disparity-selective cortical voxels. F: shows flattened cortical maps of preferred disparity of activation (see color scale bar at top) for 3 subjects, each with 3 clusters of identified voxels in cortical areas V2 (blue), V3 (green) and V3A (yellow) respectively, with projected boundaries of four visual areas (white lines: V1, V2, V3, V3A) from separate retinotopic mapping; the corresponding activity over time is shown in the inset panels, with left and right panels showing the separately tested forward and reverse conditions (see main text); the cortical clusters respond consistently to a preferred disparity for both forward and reverse conditions

Comparison of the cortical activity produced by this stimulus against a baseline activation produced by a 16 second presentation of binocularly uncorrelated set of dots revealed a set of responses in agreement with earlier studies at 3T^21^ and 7T^27^; these are shown in Figure 1C and 1D for two subjects (S1, S3) for whom repeat scans had been performed. A separate set of scans at 3-T field strength was performed to establish the functional boundaries between cortical areas V1, V2, V3, and V3A using retinotopic mapping^29,30^. The resulting assignments are shown on the anatomical slices in the right hand column of Fig 1C and the boundaries are shown as white lines and labels on the computationally flattened cortical representations in Fig 1D. The cortical activations measured at 7T are tightly confined to grey matter voxels in Fig 1C but they nonetheless show the selective activation of area V3A by this stimulus.

### Voxels show activity to a consistent disparity configuration across early visual cortex

We then searched for local topography for binocular disparity in human visual cortex by presenting subjects with planar dot patterns whose binocular depth was steadily varied with a linear ramp across the known range of sensitivity of human stereoscopic system. The ramp changed slowly over a 28 sec time course from far to near or from near to far (Fig 1B). The stimulus was divided into complementary halves. Subjects viewed a pair of planes defined by binocular disparity: one in the upper hemifield, the other in the lower hemifield. The upper plane started at a positive disparity (+0.215°), which puts the plane in the far background, distant beyond the fixation plane. Every two seconds this plane moved towards the observer passing forwards in a sequence of 14 steps, through the zero disparity plane to reach (–0.215°). When the ramp was completed, the dot patterns returned to their starting depths before the progression started again. The entire waveform therefore had a “sawtooth” pattern over time. Starting at a near disparity (-0.215°), the lower plane moved in the opposite direction to the upper plane. This configuration ensured that there was no net drive to the vergence eye movement system.

As the depth was systematically altered over the 28-second period, we recorded high-resolution BOLD fMRI images at 7-tesla field strength from the visual cortex of the subjects. Fig 1E presents a schematic of how a particular voxel or cluster of voxels that are selective for stereoscopic depth might be affected by this stimulus. As the stimulus disparity (Fig 1E: red line) changes over time, the stimulus enters the window of selectivity of the voxels (blue curve) and eventually leaves it. Each stimulus presentation projects to a BOLD response measured over time at the right, such that the peak time of response corresponds to a peak response to a particular disparity.

These data were analyzed in an analogous way to retinotopic mapping^30-32^, such that each voxel was assigned an amplitude and phase, with phase specifying the position in depth to which the voxel responded maximally. This analysis is expected to reveal the local topography for stereoscopic depth or ‘stereotopy’. All five subjects (S1, S2, S3, S4 and S5) performed this experiment at least once in the form shown in Figure 1B, E and F. We hypothesized that the opposite directions of disparity ramp should produce complementary activation patterns. Therefore, three subjects (S1, S2, S5) repeated the experiment, with the disparity shifting in the opposite direction (‘Reverse’ in Figure 1F) and two subjects (S1, S3) provided additional repeat scans with the original direction of shift (‘Repeat’).

Since we wished to maintain a high spatial resolution, the voxel size was 0.75 mm isotropic and the data were not spatially smoothed. For initial examination of the data, we generated a technique for selecting regions with spatially consistent local signals, on the basis that these voxels should have a lower standard deviation in the phase of their response in both scan sessions. This statistical measure computed the circular standard deviation (taking account of the fact that phase values are periodic modulo 2π) for the 342 voxels surrounding each voxel in native space. The circular SD computed for each voxel is a measure of the similarity of that voxel’s phase to the phases of nearby voxels, indicating local consistency across a 5 mm zone spanning the reference voxel. This procedure is explained in more detail in the Methods.

In addition to the response phase value for each voxel, there is a measure of ‘coherence’ (taking values from 0 to 1) that indicates how well the response of the voxel adheres to the sinusoidal function at the stimulus frequency relative to the response at other harmonics. This measure is used as a threshold to indicate which voxels activated by changing the disparity of the stimulus; for this initial analysis, a threshold of 0.35 was used. Applying the analyses described above, only voxels that had circular variability of less than 2 standard deviations of response phase were considered for subsequent analysis, with each remaining voxel then assigned a phase value indicating the preferred position in depth.

Voxels passing the clustering procedure and exceeding a coherence value of 0.35 are shown in Figure 1F, color coded according to the phase of the response indicated in the rainbow-colored bar at the top each column. Specific regions of interest in V2 (blue box), V3 (green box), and V3A (yellow box) were chosen for the ‘Forwards’ direction data for each subject. The phase preference of these voxels reversed when the direction of the planes was reversed, indicating that the response is related to the stereoscopic depth of the plane. The BOLD response to the average disparity cycle for sample voxel clusters in V2, V3 and V3A for the three subjects with both ‘Forward’ and ‘Reverse’ scan sessions is shown in Fig 1F (inset graphs), with the phase shift between the two stimulus directions being evident in the plots. Specific clusters within the visual areas were chosen on the basis that they (i) were ‘active’ in both directions of stimulus progression i.e. had a coherence above 0.35, (ii) were entirely within a defined ROI (V2, V3 or V3A) and (iii) consisted of a minimum of 20 voxels. The coherence value of 0.35 was chosen as it reduced the number of clusters available. We considered clusters of voxels, because the 5 month time lapse between scan sessions and small discrepancies in slice prescription and registration, precluded exact alignment of individual voxels across sessions.

Fig 1F shows that size of the BOLD signal changes are large, on the order of 1-2%, indicating that the stimuli generate strong responses in these early visual areas. In order to determine the depth configuration to which the voxels were most responsive, a hemodynamic lag of 4 s was assumed. With this shift applied, it is clear that the responses are to different locations in depth, not just the time point at which the ‘jump’ between near and far occurs. In Subject 1, the preferred disparity of the cluster from dorsal V2 is when the lower field of dots is ‘behind’ fixation – the peak is around 6 sec, so assuming a hemodynamic response of 4 seconds, this would be the 1^st^ stimulus position (±0.18°). In contrast, the peak for the reverse phase is around 22 sec, giving a maximum response when the lower field is once again ‘behind’ fixation. We can also reject the possibility that the responses are dominated by similarity of depth in upper and lower fields (i.e. a preference for zero disparity), as this would result in a peak response at around 18 s in both stimulus directions as the two planes are closest to zero disparity at the mid-point of the cycle for both ‘Forward’ and ‘Reverse’ conditions. While there are examples of a peak at this point, there does not appear to be over-representation of this response phase.

Whilst this clustering procedure succeeded in detecting clumps of cortical activity that are convincingly related to the stereoscopic depth content of the stimulus waveform, the procedure requires some local spatial combination of nearby voxels to reveal the clusters and a degree of choice based on human inspection of the data field. Both of these limitations mean that the estimation of cluster size is to some extent dependent on the analysis procedures themselves. So although we can conclude from this analysis that there are clustered responses to stereoscopic depth in extrastriate cortex, we cannot firmly conclude the size and shape of these clusters and we cannot deduce anything about the proportions of cortex occupied by high signal clusters.

### Variogram analysis of disparity clustering

The resolution of these 7T functional images is 0.75mm in linear dimensions, while the columnar structures that we seek may be anything from 2-10 mm in size across the cortical surface. Our aim is to find a link between spatial position on the cortical sheet and the functional properties of neurons, so it is essential to dissect apart the different sources of covariation between BOLD signal and spatial position on the cortical surface. Fig 2A highlights the issue with a sample of cortical activity from cortical area V3A of subject S5: the pattern of cortical activity is clumpy in its spatial distribution. We shall show here that the size of these clumps is signal-dependent.

**Figure 2:**
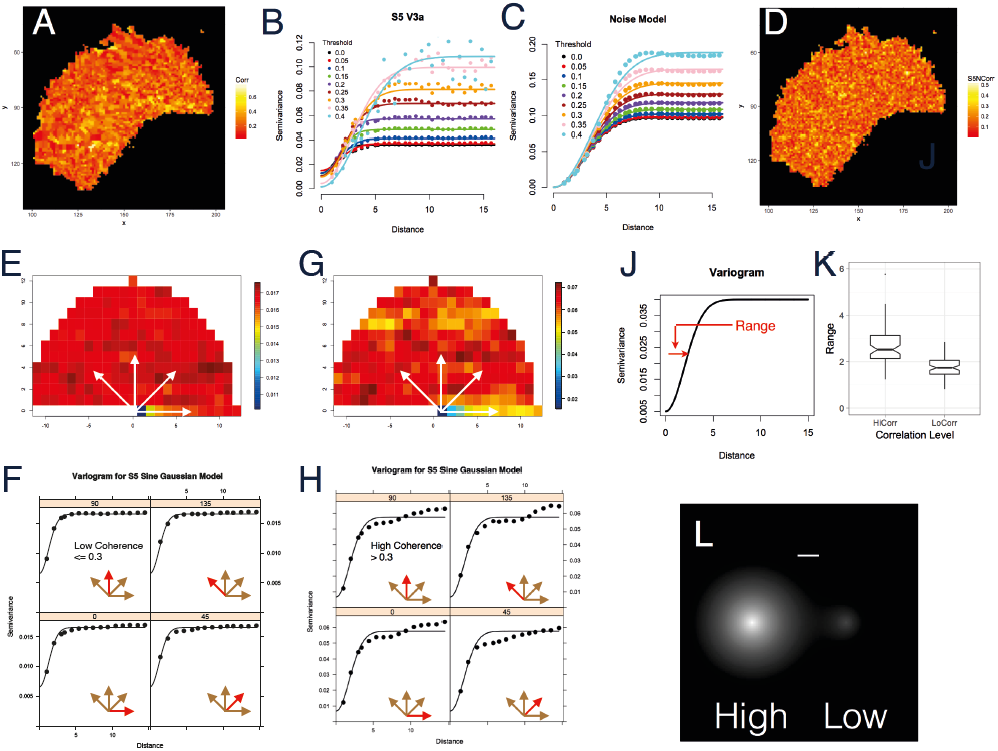
Characterizing the spatial correlations of BOLD ; data. A: shows a cortical flat-map of a field of stimulus-related BOLD activations (Subject S5, area V3A) to highlight the clumpiness of these responses: color-coding shows the amplitude of the coherence (Corr) between time-varying BOLD and the stimulus. B: variogram curves showing variance between neighboring samples as function of cortical distance on flat-map from data in Fig 2A thresholded by coherence level; the variance between nearby spatial samples is lower than the variance between more distant samples, reflecting local spatial correlations in the data; as threshold is increased, these spatial correlations become more long-range for high coherence signals. C: variograms from simulations of spatially correlated noise similar to the low coherence signals from subject S5. D: simulation of cortical flat-map of correlated noise for subject S5 area V3A using fitted parameters for low coherence signal (< 0.3 correlation). E: Variogram of low coherence signals (<=0.3) from subject S5 as a function of distance and direction across cortex; data for directions along white arrows plotted in Fig2F: variograms in four cardinal directions. F: variograms of the sine component of coherent cortical activity for subject S5 for low coherence data measured in four cardinal directions (0, 45, 90, 135); a cumulative Gaussian curve is fit to the results and the standard deviation measures the range of spatial correlations). G: Variogram of high coherence data from S5 shows evidence of longer-range spatial structure compared with E. H: Variograms of high coherence data to compare with Fig2F, poorer fits to the data and deviations from the fitted Gaussian curve indicate some spatial anisotropy for high coherence signals. J: the Gaussian curve, particularly the range parameter (red), defines local spatial correlations of the BOLD signal. K: box plot showing difference in fitted range parameters between high coherence (>0.3) and low coherence (<=0.3) responses across four cortical areas (V1, V2, V3, V3A) for all subjects. L: comparison of cortical spread for high coherence signals (left) and low coherence (right): the high coherence spread has a wider range parameter as in J and a 4x higher amplitude (compare F and H); scale bar (top) shows 7.5mm (10 pixels).

There are a number of structural and functional factors that lead to incidental, local covariation of the MRI signal across the cortical surface. Some of these are due to the imaging method itself, others are due to the local anatomy of blood vessels or cortical folding, whilst some local covariation will be genuinely related to neural function. To test claims about the specific form of covariation that is implied by a columnar organization for a functional property requires separate identification of the signals relating to cortical columns from the factors that are incidental. This circumstance often arises in other areas of biology and earth science, particularly in data arising from fieldwork.

To quantify the clustering of disparity, and particularly to reveal how the spatial covariation of disparity preference differs from correlated noise, we adopted an approach that is common in environmental modeling of various kinds^33^ but little used in neuroimaging studies (for exceptions see Ye et al^34^ and Bellec et al^35^). If we characterize the variance of signals within a voxel inthe usual way *Var(X)* =*E(X*^*2*^*)-E*^*2*^ *(X),* then the variogram captures the spatial dependence of variance by parameterizing the distance between ents *Z(x, y)* at points separated *by(dx,dy),* as follows:

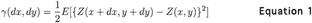

where *Z(x, y)* is the activation at point *(x, y)* and E[] is the statistical expectation operator. Note that the quantity γ (sometimes called semivariance because of the factor of 1/2) depends only on the separation of points (*dx, dy*) and not the actual locations of the points (*x, y*). The variogram is a statistic based on the distribution of individual point samples, rather than a continuous noise model.

Fig 2J shows a typical outcome from this analysis. The value of γ increases with distance, demonstrating that nearby values are more closely related to each other than more distant values. Eventually further increases in the distance parameter make little difference. The curve is a cumulative Gaussian, whose standard deviation parameter, here termed the “range”, gives a metric, in pixel units, for the distance over which the local similarity of nearby values persists. If the data contained no spatial correlations, the variogram would be flat. If the data field were continuous rather than a set of discrete sample points, then the local spatial correlations could be described by the spatial autocorrelation function: for statistically homogeneous data, there would be a direct relationship between the range of the calculated variogram and the spread of the autocorrelation function.

In the current dataset, the data of particular interest are those that have high coherence with the changing stimulus waveform, meaning that the response of those voxels is significantly related to the disparity-defined stimulus. Fig 2B shows the variogram for the data field in Fig 2A at a succession of increasing threshold values for the coherence level. All values below the threshold are discarded. At low values of threshold, the variogram rises and then remains flat. At higher values, the location of the asymptotic point rises, reflecting the fact that the threshold has selected data with larger values of coherence. At the largest thresholds, the curve pushes to the right along the x-axis. This indicates that the average distance over which the data remain locally correlated increases with larger signal strength. Fig 2C and Fig 2D present the equivalent calculations for a random field of spatially correlated noise. The spatial correlations in the noise are closely matched in Gaussian form to the correlations in the low signal components of the experimental data.

Given that the experimental curves show a different spatial form for higher coherences near to the value of 0.3, for the rest of this analysis, we take the split between low and high coherence signals at a threshold of 0.3. We note that this choice of threshold has been typical for other MRI analyses using periodic stimuli, as in retinotopic mapping, but we found that the exact value chosen did not substantially affect the pattern of results.

Because the variogram depends on the separation of points (*dx, dy*), it is possible to define angular sectors relative to each starting point (*x, y*), which introduces an anisotropy term into the specification of the variogram (see Figs 2E, 2F, 2G, 2H). Fig 2E show examples of variograms for subject S5 for low (< 0.3) coherence, whilst Fig 2F shows the same for high (>= 0.3) coherence. Fig 2E (low coherence) also presents the calculation of the variogram in four cardinal sectors relative to the reference point. There is little evidence of any anisotropy at low coherences values and indeed the variogram is well described by the cumulative Gaussian fit at larger distances. Fig 2F shows that the range is significantly greater in the high coherence condition. Although the cumulative Gaussian describes these data quite well at small interaction distances, there is some evidence of a poorer fit for the cumulative Gaussian curves and some evidence of anisotropy, both particularly apparent in the deviations from the estimated Gaussian at larger interaction distances. For the most part, the assumption of isotropy (radial symmetry) is adequate for the low coherence (< 0.3) signals (see Fig 2E and 2F). Our aim here is to create a suitable empirical model of the low coherence spatial correlations as a basis for simulation-based statistical testing of the spatial structure of the high coherence signals, specifically searching for evidence of clustering or columnar structure. Therefore, we adopted a modeling framework based on the assumption of isotropy, even though there is some evidence that this assumption is violated for high coherence signals.

Using the assumption of radial symmetry, we calculated the variogram on an individual basis for data from each subject, stimulation condition, cortical area, and coherence level (low vs high) under the assumption of radial isotropy. A box-plot showing the distribution of the range parameter from this analysis is shown in Fig 2K. This confirms the impression from the individual data in Fig 2F and 2H that fits of the cumulative Gaussian to high coherence stimulation conditions lead to larger estimates of the range parameter. We applied an ANOVA to this full set of fitted range values, specifically to compare the significance of any difference between high and low coherence conditions. A simple ANOVA comparing the ranges for low versus high across all conditions is highly significant (F(1, 149) = 83.2; p < 4.7E-16), as is the main effect of low vs high coherence in an all-ways comparison of coherence level, subject identity, and cortical area identity (F(1,111) = 93.7; p < 2.2E-16). There is a weaker effect of subject identity (F(4, 111) = 5.56; p < 0.0005) but no main effect of cortical area or any significant interaction effects. A main effect of cortical area or an interaction between coherence level and cortical area might have been predicted on the grounds of the different vascular structure of V1 in comparison with other extrastriate visual areas, but nothing is discernable at this scale of measurement.

The spatial correlation structure of the low coherence data represents a practical limit on the resolving power of the imaging methodology. Data samples that are partially correlated with one another provide less information about the underlying signal than data samples that are independent. Our analysis methods have stripped away most of the elements from a conventional MRI analysis pipeline that would introduce spatial smoothing across the data points, with the exception of motion-correction for head movements. Motion and errors in its computational correction will appear as long-range effects in the variogram curve, raising the variance equally across all distances of the variogram curve: in the language of environmental modeling, this will increase sill variance but is not expected to affect the range. A second methodological concern is whether geometric distortions introduced by the computational unfolding of 3D scanning data onto a flat-map representation of neural activity might affect the estimates of spatial correlation. It is hard to see how the unfolding procedure could generate a spurious difference between voxels with low and high coherence signals, since these voxels are so tightly intermingled with one another. As a further check, we performed the entire variogram modeling procedure in 3D native imaging space (see Supplementary Fig 1); the difference of spatial correlation distance between low and high coherence voxels remained highly significant (F(1,126) = 17.025; P < 6.654E-05).

Figure 2K therefore indicates that there is a significantly greater clustering in the high coherence data across all visual areas. Greater clustering at high coherence means that these stimulus-related activations tend to occur in disparity-specific clumps of responding voxels. In order to visualize the difference in the sizes and shapes of the clumps between the low and high coherence data, a summary distribution is shown in Fig 2L. These are radially symmetric Gaussian distributions with a standard deviation taken from the fitted range parameters in Fig 2K. The peak height of the two Gaussian functions also differs, being a factor of 4 higher for the high coherence case, which is similar to the difference illustrated in Fig 2 G and H (compare vertical y-axis scales). We conclude that the high-coherence data, which is the signal-related component of the BOLD response, resides in clumps that are larger and more extensive across the cortical surface than the low-coherence data, which may be regarded as spatially-correlated noise. This possibility has been previously acknowledged^36^.

### Local spatial correlations intrude upon the identification of stimulus-related clusters

The presence of local spatial correlations both in the low and high coherence signals presents a challenge for the reliable identification of clustered or columnar structures on the cortical surface. In the animal imaging literature, computational models of the neural development of cortical columns often depend upon local spatial correlation of neuronal signals combined with a pattern-inducing threshold step^37,38^. To the extent that the computational analysis of MRI data includes these essential, mathematical components, clustered or columnar structures may also emerge, consequent upon the presence of local spatial correlations in the data that are passed through a non-linear threshold or classification.

We used the fitted estimates of the cumulative Gaussian curves for each variogram (as shown in Fig 2F) to estimate the baseline level of spatial correlations in the low coherence data as a practical estimate of the interfering noise against which the high coherence signals must be detected. By noise here, we include all types of interfering components: true noise as well as low level signals similar in structure and sources as the signals that we want to isolate and measure. The resulting noisy patterns (Fig 2D) have local spatial correlations between neighboring pixels that match the local spatial correlations found within the low coherence signals of the experimental data, but each new sample pattern that is generated is statistically independent of the other patterns.

The noise simulations yield amplitude and phase components, similar to the experimental data. The simulated spatial distribution of phases was plotted across a cortical area as in Fig 3B and 3D for comparison with the experimental data (Figs 3A and 3C). These data plots and simulations provide a color-coded representation of phase (rainbow map, as in Fig 1F) for each pixel within the cortical areas V1 (Fig 3A and 3B) and V3A (Fig 3C and 3D) from subject S5, including both high and low signal coherences. The distribution of phases in the noise simulations has local similarities, which are generated entirely by the local spatial correlations. With appropriate choices of parameters, the local similarities of phase predicted by the simulations can become strikingly similar to the patterns of columnar structures reported in the cat and monkey visual cortex (see Supplementary Figure 3). Nonetheless the experimental data shown in Fig 3A and 3B are distinct from the simulations in Fig 3C and 3D. Particularly for area V3A, the data show larger sized regions with a consistent representation of stereoscopic depth: this is evident in the larger size of zones with a single color in Fig 3C (V3A), compared with Fig 3D. In V1, the data show evidence of a clustering of preference for similar stereoscopic depth along the line of the horizontal meridian (Fig 3A, V1). The main difference between the spatial structure of simulations in comparison with the experimental data is that the simulations in Fig 3B and 3D are constructed to reflect only the spatial correlation structure of the low coherence signals, whilst the data shown in Fig 3A and 3C contain both low and high coherence signals.

**Figure 3:**
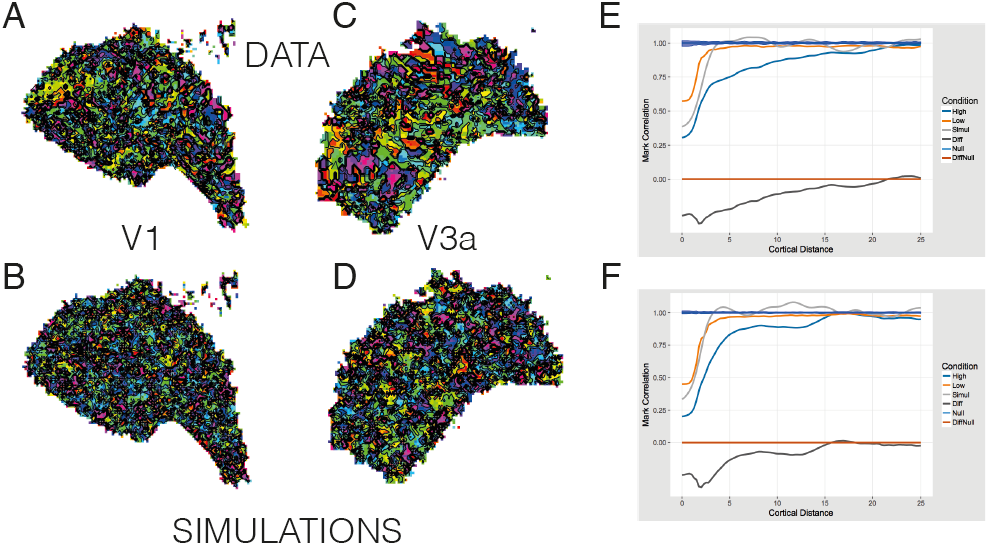
Low coherence responses give baseline level of spatially correlated noise for testing the spatial distribution of responses in experimental data. Colored regions are larger in extent in the experimental data, especially in V3A, showing consistent responses to single disparities. A and C: Experimental values of response phase for all signal strengths, for subject S5, cortical areas V1 and V3A, color coded as in Fig1F. (The streak of consistent green/turquoise response phase in the V1 experimental data is most likely due to the response of the horizontal meridian.) B and D: Simulations from synthesized noise of preferred disparity of pixels across the cortical map: rainbow color-coded contour maps of response phase for cortical areas (V1, V3A) from subject S5. E: mark correlation function plots correlation between phase values for pairs of voxels as function of spatial separation, Subject S1, area V3A. High: V3A data for coherence > 0.3; Low: V3A data for coherence <= 0.3; Simul: synthesized noise (as in Fig 3 D) matching spatial correlations of low coherence data; Diff: difference between High and Low curves; Null: predicted value of mark correlation with random reassignment of phases at each voxel, blue band shows estimated 95% confidence based on point wise resampling. DiffNull: predicted value of Diff for no difference between High and Low. F: as E but for subject S3, area V3A. Spatial extent of Diff curve indicates structured representation of disparity across cortical surface over distances as great as 11-15mm.

Further quantification of the spatial similarity of preferred phase between nearby pixels on the cortical flat-maps can be achieved with the mark correlation function employed in spatial st at istics^39^

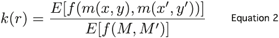

where

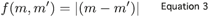

Figs 3E and 3F plot the mark correlation function for phase angle as a function of distance across the cortical surface within cortical area V3A for subjects S3 and S1 respectively. The turquoise curve plots the mark correlation for the high signal pixels (stimulus coherence >= 0.3): for both subjects, this curve stretches out to a distance of 15 or more pixel elements. This measure of interaction distance can be considered as equivalent to the range parameter introduced earlier in the paper. The orange curve plots the mark correlation for the low signal pixels (stimulus coherence < 0.3); this curve is more restricted in spatial extent, reaching a value close to 1 at distances of 5 pixel elements. The light grey curve is based on the computational simulation of the low coherence data; it has a similar spatial range. The dark grey curve (Diff) is the difference between the high and low coherence mark correlation curves; it is expected to be zero where the two correlation curves are the same and it highlights the greater range of interaction distance for the high coherence pixels. The null curve represents the null hypothesis of random association between marks and spatial positions and the bootstrapped limits around it represent the pointwise estimates of 95% range based on random resampling. The diffnull curve is simply a reference line at a value of zero.

These calculations confirm the greater range of spatial correlation of marks in the high coherence data, suggesting spatial correlation of phase values out to 15-20 pixels, equivalent to 12-15mm across the cortical surface. Further examples of mark correlation curves and the distribution of preferred phases across four cortical areas (V1, V2, V3 and V3A) for subjects S3 and S5 are presented in Supplementary Figures 5, 6, 7,and 8. These data confirm the large range for spatial correlation of preferred stereoscopic depths in V3A and other extrastriate cortical areas. The analysis also indicates a tighter range for the spatial correlation of high coherence signals in cortical area V1.

### Repeated measures of spatial structure in responses to stereoscopic depth

One way of addressing the challenge presented by structures created by local noise correlations is to seek repeated measurements, as in Figure 1F. There are two reasons for seeking alternatives in addition to this approach. The first is that non-invasive MR imaging data is often inhomogeneous. There are often consistent causes of signal dropout, such as large blood vessels or partial voluming effects at the boundary between cortical white and grey matter. Taking repeated samples does not help with this source of signal variation. The second issue is practical: MR imaging experiments are often quite lengthy and it is often difficult to achieve multiple repeated imaging sessions with volunteer subjects, so methods that could be applied to data from a single imaging session are potentially valuable. Nonetheless, the statistical methods we have introduced here ought to provide consistent outcomes under repeated measures.

Figure 4 presents the outcome of repeated measurements of the disparity phase-mapping procedure on three occasions with subject S1. The measurements took place at intervals spread over several months. Two of the data sets are with identical stimulus sequences (‘forward’ and ‘repeat’), whilst the other (‘reverse’) presents the same disparity waveforms but traversing the stereoscopic depth range in the opposite direction. Figs 4A, 4B, 4C show the measurements of preferred phase across cortical area V3A for subject S1, as in Fig 3C for subject S3. These experimental data include both high and low coherence signals. The maps are color-coded as before and the black lines distributed across the map represent equal phase contours, with the density of these lines showing the local gradient of the phase landscape. Figs 4D, 4E, and 4F result from the differentiation of the maps in Figs 4A, 4B, and 4C: the differential of a 2-D scalar variable is a vector quantity with amplitude and direction (corresponding on a landscape to the size of gradient and the direction of steepest descent). In Figs 4D, 4E, 4F, the amplitude of the directional derivative is color-coded such that green represents flat regions, where the disparity phase in the corresponding figures above (Figs 4A, 4B, 4C) changes only slightly from one pixel to its spatial neighbors, whilst regions with large local changes in disparity phase show as yellow or orange. The contoured surfaces in Figs 4D, 4E, 4F are therefore a differential landscape map of the disparity responses across the cortical surface.

**Figure 4:**
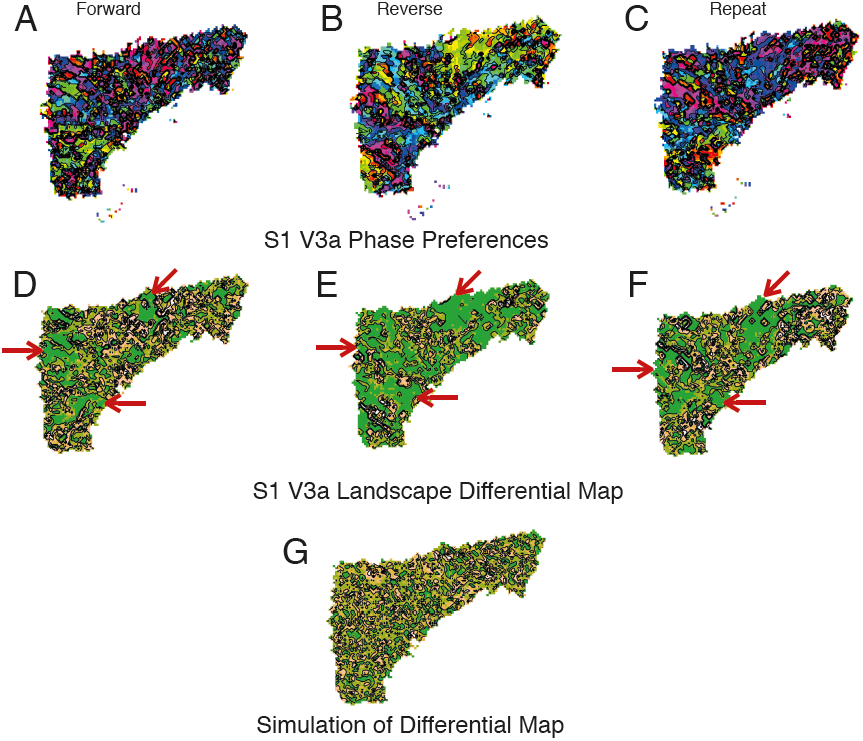
Extended regions of V3A respond consistently to binocular depths. A: Experimental values of response phase for all signal strengths, for subject S1, cortical area V3A, color-coded as in Fig1F for Forward movement of disparity planes. B: similar to A, but for reverse movements; preferred phases change. C: Repeat scan for Forward movement, as in A. D: differential map of amplitude of phase changes across cortical surface; topography scale of amplitude of phase differences, from green (small difference) though light green to yellow to white (large difference); density of contour lines (black) shows local steepness of gradient. E: similar to D for reverse movement as in B. F: similar to D, E for repeat measure. G: differential landscape map calculated for synthesized noise with spatial correlations matching low signal voxels. In all cases, green area of data maps are larger than synthesized noise demonstrating local similarity of response to disparity phase. Red arrows in D, E, F highlight similar features of differential landscape map across repeated measurements.

Conversion of the phase preferences to a differential landscape map facilitates the comparison of the different stimulus conditions. The low amplitude regions of the differential map are expected to be similar across forward, reverse and repeat conditions and the red arrows in Figs 4D, 4E, 4F highlight areas of consistency between these three separate scans. These areas of consistency imply that in these regions there is a high degree of similarity of the phase response, across both low and high coherence signals. Fig 4G shows the differential landscape map for the simulated data based on spatial parameters that match the low coherence signals measured within the experimental data.

Figure 5 puts together, for comparison, experimental data and simulations extracted from the remaining 4 subjects tested in this study for area V3A. In every case, the experimental data show much more extensive regions, whose differential has low amplitude (green), in comparison with the simulations based on low-coherence signals. This shows that V3A always has extended regions where there is a consistent representation of stereoscopic depth.

**Figure 5.**
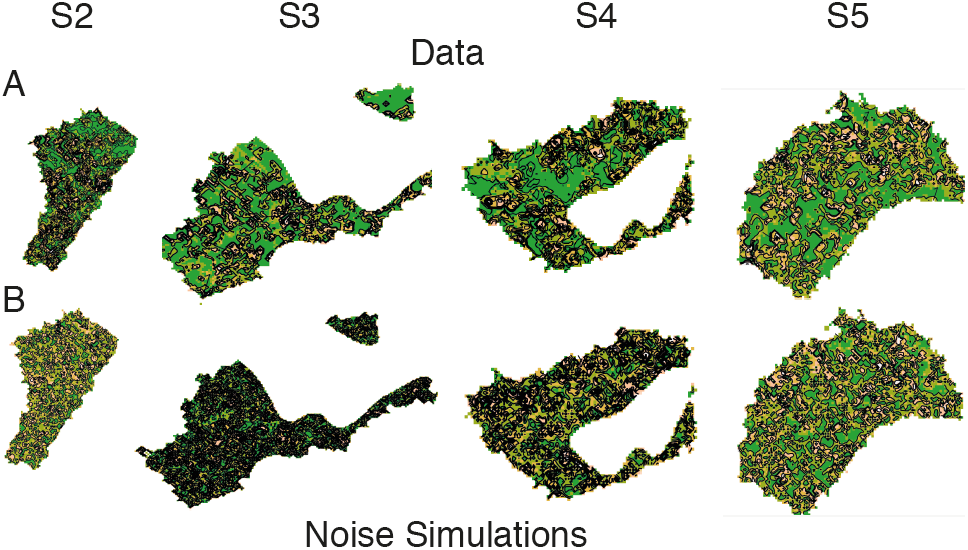
Comparison of differential maps of experimental data (A) for cortical area V3A against the expected differential maps from noise simulations (B) for the remaining four subjects tested (S2, S3, S4, and S5). All experimental data consistently show greater extents of responsiveness to similar disparities—green regions are larger in experimental data than in simulations, color coding and contour lines as in Fig 4D, 4E and 4F.

In summary, we show that spatial correlation analysis can distinguish the spatial organization of high coherence signals for disparity from the spatial organization of low coherence signals. We use this analysis to demonstrate that the high coherence signals for stereoscopic depth in cortical area V3A are not simply thresholded out of a noisy background containing local spatial correlations. The high coherence signals are clustered together to form patches of cortex with similar disparity preferences. We also show in Figs 4 and 5 that V3A contains spatially extensive regions of cortex, across which the preferred disparity is quite similar from one point to the next. The size and organization of these spatially extended regions are too large to be compatible with the spatial correlation structure of the low coherence signals. Analysis by means of the mark correlation function suggests correlated structure out to 12-15mm across the cortex. Although these features of cortical mapping are defined statistically, they are present in all subjects tested and in repeat testing of individual subjects.

### Representation of stereoscopic depth in cortical area V3A

The preceding analysis showed that adjacent voxels in cortical area V3A tend to have similar preferences for disparity and that high coherence signals are present in larger than average clusters. Results from monkey imaging experiments show evidence for a progression of disparity across the cortex, most notably in area V2^14^. Given the evidence for specialization of function in V3A, we wondered whether a similar progression of preferred disparity could be demonstrated for this area in human visual cortex.

As previously, we compared the phase-encoded, high coherence data to a distribution of phases derived from computational simulations of spatially correlated noise matching the characteristics of low coherence data. We performed two different analyses to determine whether disparity phase varied systematically across the cortical surface. In the first of these, the analysis of phase progression proceeded by selecting cardinal directions (0, 45, 90, 135 degrees) within each cortical area. Along lines parallel to these cardinal directions, the phase values were unwound, differentiated, and then subjected to a run-length analysis to identify consistent regions with succession of phase advance or phase lag.

We compiled frequency histograms of the distribution of run-lengths, taking only run-lengths greater than 2, for the noise samples and true data. We compared these distributions to determine whether the progression of disparity preference across the cortical surface differs from that found in spatially correlated noise.

Whilst the synthesized noise samples are continuous, the actual data at high-coherence levels are discontinuous because there are regions where the signal coherence drops below the threshold level (0.3). Notwithstanding this point, the true data showed a greater tendency to contain longer run-lengths of consistent change of preferred disparity. Of course, the spatial correlations present in the matched noise samples mean they have greater run lengths of phase than noise with no spatial correlations. But the experimental high coherence data show an even greater number of cases of long run-lengths than the matched noise samples (Kolmogorov-Smirnov test: see Methods for details). An analysis of the proportion of run lengths greater than 3 voxels indicates that, while this value does not differ between V1, V2, and V3, V3A shows a significantly greater proportion of these longer run lengths.

Comparison of these long run-lengths with the phase maps of the type shown in Figs 3 and 4 revealed the source of these extended zones. They tended to lie on the slopes of the phase-landscape, traversing the side of a hill or valley. Therefore the presence of these long run-lengths does not seem to be similar to the progression of orientation preferences found in the cortical map of V1. Rather they arise because the spatial extent of hills and valleys in the landscape of disparity-phase responses is greater and this greater spatial extent arises from the clustering of high signal regions identified in the previous section.

As a more refined check on whether there is statistical evidence for a progression of preferred phases, we employed the mark correlation function used in Fig 3E, 3F. In this case, however, we took the distribution of marks as being the vector directions derived from the differential landscape maps. An example of the arrow-map, showing local directions of phase change, superimposed on the differential landscape map, showing local amplitude of gradients, is shown in the upper panel Supplementary Figure 9 for area V3A, subject S5, reverse condition, together with the mark correlation function for the vector directions. There is a slight tendency for phase differences to be dissimilar at a distance of 3-4 sampling units (2.3-3.0mm). This is consistent with the conclusion from the phase run-length that there are changes of preferred disparity phase on that scale. To proceed further would require better segregation of the distribution of true stimulus-related signals from the structured background of neural dust. This will need both better data so that both signal and background can be more accurately characterized and further development of the statistical model.

## Discussion

Our search for spatial structure for stereoscopic disparity within visual cortical areas focuses on several key principles. First, we have searched for physiologically relevant aspects of the BOLD signal: our use of high spatial resolution imaging at 7T allows the cortical activation to disparity-defined stimuli to be clearly attributed to the grey matter of the cortex, so the disparity-related component of the BOLD responses can be readily identified. These BOLD responses change smoothly and systematically as the disparity of the stimulus is changed. Second, we have concentrated on those elements of the imaging data that can be demonstrated in all subjects tested and in repeat tests of the same individual. Third, previous explorations of columnar structure have sought to classify parts of the cortex as dominated by stimulus class “A” in comparison with stimulus class “B”. We show that in 7T MRI images that this A vs B comparison is vulnerable to variation in the signal quality and also the spatial correlation structure of background noise.

We therefore employed a combination of continuous variation of the stimulus values, together with quantitative descriptive statistics to separate the low and high signal regions of the cortex. We find that disparity-specific cortical responses are clustered within domains of the cortical map, within which nearby points have similar preferences for the stereoscopic disparity of the stimulus.

### Organization of cortical representation of stereoscopic disparity in area V3A

Previous studies have built a strong case for cortical clustering of selectivity for visual attributes such as orientation and ocular dominance. By contrast, little is known of the organization for stereoscopic disparity in the human early visual cortex, despite the fact that there are some long-standing proposals for a disparity-based ordering in mammalian cortex^40^. Based on the literature from the macaque monkey, V2^14,15^, V3^16^ and V3A^16^ have a clustering of disparity selective neurons, and therefore could be expected to show clusters of voxels preferentially activated by a disparity-defined stimulus. Perhaps not surprisingly and consistent with recent work in human cortex^27^, we found that V3A shows the strongest evidence for cortical domains for processing stereoscopic depth, likely resulting from an ordered connectivity of disparity-selective neurons. We find that the prominent characteristics of V3A for disparity processing are the presence of strong disparity-related modulations of BOLD activity and a broad range of stimulus preferences for stereoscopic depth. It is for this reason that the original mapping stimulus used by Tsao et al^21^ to highlight V3A is so effective, because it relies upon a comparison of activation to a range of depth planes against the activation to a single depth plane.

The topographies we identify here are presumably largely driven by post-natal experience-dependent processes40 and their size and shape within the cortical surface may reasonably be expected to vary in detail from one individual to another, dependent on visual experience. Nonetheless, there are consistent features of these disparity-specific regions, which are the local consistency of disparity preference within a region (Fig 3) that is also present when the stimuli are presented to the same individual in the opposite direction (‘reverse’).

The distance over which interactions are generated in these measurements is of the order of 12-15 mm of cortical separation. This distance is within the range of a combination of monosynaptic and disynaptic cortical intrinsic connections, specifically horizontal intra-areal connections in area V5/MT of extrastriate cortex^41^. It is also possible that feedback connections from one cortical area to another may contribute to the functional interactions we visualize here^42^. We suggest that the functionally defined regions we have discovered here will ultimately be proven to align themselves with anatomical markers such as Cat 301 staining^43^, which are associated with disparity processing in non-human species.

### Potential impact of vergence eye movements

A chief reason for splitting the upper and lower visual fields and changing the disparity in opposite directions was to prevent subjects dynamically adjusting convergence to the plane of the dots. By having fixation at zero disparity and the planes symmetrically arranged in depth around that point, the drive for vergence is considerably reduced. Since it was not possible to monitor eye position in the scanner, subjects were recruited who were experienced at viewing visual stimuli, and known to be able to maintain fixation. As to whether the results could be explained by vergence eye movements, the primary impact of vergence would be on neurons that have sensitivity to absolute disparity, since changes of vergence change the whole pattern of absolute disparities across the visual field whilst leaving the relative disparity between visual features unchanged^12^. Any effect of vergence in altering the presumed stimulus disparity at the retina would have to be consistent across all voxels with sensitivity to absolute disparity. Thus, if vergence did have an impact on cortical activations, this should be more evident in V1 than extrastriate cortex. By contrast, our results show a structured pattern of cortical responses that are stronger in extrastriate visual areas than in V1.

### Voxels are not just responding to large changes in depth or to over-representation of the zero disparity (the ‘horopter’)

A potential explanation for the clustering of disparity preference could be the response of the neurons to the large change in disparity: that is, the ‘jump’ from near to far or far to near at the end of the cycle. However, if this were the case, the preferred phase of the response should be consistent throughout the visual areas and conditions, as the step change occurs at the same phase in both halves of the visual field and in both directions of movement in depth.

Alternatively, if there were a large representation of the horopter, the voxels should show tuning for a phase corresponding to the point at which the disparity difference between the upper and lower planes is minimal. In both the ‘Forward’ and the ‘Reverse’ condition, the point of maximum stimulation of the horopter occurs at the middle phase. Thus, the peak would be predicted to arise at time points around 18-20 s in both conditions. We see evidence for such a structured organization along the horizontal meridian in visual area V1 (Fig 3A). In other areas of early visual cortex, while there are a few regions where the peaks align for the two conditions, for the majority of activated voxels in V3A, there is a phase difference between the two conditions.

### Clusters in the field of fMRI responses

Our results and analytical methods also contribute to the problematic analysis of clusters in functional MRI data^6^. We show that high signal regions are grouped into clusters, that these clusters are of a different size than low signal regions, that these high signal clusters are most likely spatially anisotropic in shape and that these high signal regions convey physiologically meaningful information about the stimulus when the stimulus waveform is altered. We also find that the spatial variogram of these high signal regions may have deviations from a simple Gaussian model at longer distance interactions. Our current thinking is that the deviation we find is related to the long-tails deviation of signal distributions noted elsewhere in MRI data^6^. More extensive data sets will be need to characterize the statistics of the tails of high signal distributions.

## Conclusion

This study has shown that there are clusters of disparity sensitive neurons in V2, V3, and V3A. Furthermore, within V3A there are clusters of voxels tuned to specific disparities and organized into cortical domains, which show a consistent appearance in spatial structure across different experimental conditions and repeated scanning sessions. An important next step is to determine whether this clustering is related to other functional organization, such as color or orientation. The geostatistical analysis of the properties of cortical responses provides a secure basis for the quantitative measurement of many properties of cortical maps.

## Methods

### Subjects

Five subjects with previous experience of fMRI experiments and normal stereoscopic vision (as determined using the TNO test) participated in the study (age range 22-37; 4 females; 1 male). All procedures were approved by the Medical School Research Ethics Committee at the University of Nottingham, with written consent obtained from all subjects. All subjects participated in a retinotopic mapping session at 3T to delineate boundaries between visual areas in the occipital cortex and allow assignment of functional activation to specific areas. The two main experiments (see Figure 1A and B) were performed at 7T; two subjects performed Experiment 1 and Experiment 2 once each (Subjects 2 and 4); two subjects performed Experiment 1 twice and Experiment 2 three times, 5 and 24 months apart (Subjects 1 and 3). Subject 5 performed Experiment 2 twice in the same session. When Experiment 2 was performed twice, the planes changed in opposite directions of depth over time.

### Stimuli and experimental design

The experiments were designed to investigate activation to disparity signals in the early visual areas with high spatial resolution fMRI at 7 T. Viewed monocularly, the stimuli consisted of patches of light and dark red and green randomly positioned dots, to which binocular disparity could be applied. All stimuli were presented using a projector, viewed by the subject on a screen at their feet through red/green anaglyph prism glasses and an angled mirror. The viewing distance was 240 cm, and the stimuli subtended a visual angle of 10.75°.

In Experiment 1, random dot stereograms consisting of 200 light and 200 dark dots (dot size = 0.12°) were presented on a mid-contrast background. The central region was divided into a grid of 3 × 3 squares. The central square contained a fixation cross and remained at zero disparity throughout the ‘disparity’ block. The disparity of the 8 surrounding squares changed every second to a disparity randomly selected from ±0.1°, 0.2 °, 0.4° or 0°. A schematic diagram of the stimulus is shown in Figure 1A. This ‘disparity’ block was presented for 16 s, alternating with 16 s of dots uncorrelated between the two eyes. A single scan run consisted of 6 repeats, totaling 48 volumes. Two of the subjects were scanned twice to test for reproducibility.

In Experiment 2, the stimulus was divided into upper and lower visual fields and equal and opposite disparity was added to the planes. At the start of a cycle of the experiment, the upper plane contained a positive disparity such that it was behind of the fixation plane (Figure 1B). Every 2 s the plane moved away from the observer by a disparity step of 0.033°, passing through the fixation plane to a negative disparity of -0.215°. The lower plane did the opposite, starting at a near disparity plane of -0.215° and moved to +0.215° in counter-phase with the upper plane. At the end of the 28 s cycle, the upper and lower planes jumped back to the starting disparity values and the cycle began again. This condition is referred to as ‘Forward’. Each scan consisted of 6 cycles, giving a total scan length of 168 s. There were 8 scans in each session. This session was repeated in two of the subjects. Three of the five subjects were scanned in an additional scan session with a similar stimulus. In the second session the upper plane began at a near disparity (-0.215°) at the start of the cycle and moved backwards in depth to +0.215°, while the lower plane started at far disparity and moved forwards (the reverse of session one), termed ‘Reverse’.

Retinotopic mapping data (TR = 3 seconds, 2 mm isotropic resolution, 46 slices) and a high-resolution T1-weighted whole head anatomical image (MPRAGE, 1 mm isotropic resolution) were acquired on a Siemens Verio 3 tesla whole body MRI scanner (FMRIB Centre, University of Oxford) using a 32-channel receive coil. Stimuli were viewed on an LCD monitor at a viewing distance of 216 cm, giving a visual angle of 8° eccentricity. The stimulus presented was a luminance wedge, or a 45° black and white checkerboard wedge on a grey background rotating counter-clockwise around a red central fixation point, with a flicker rate of 8 Hz. The wedge shifted position in its rotation every 3 s, with a total of 12 positions in one full cycle. Each run took under 4 minutes (6 cycles/run), with a total of 6 runs in one session for each subject, except for subject 2 who completed 4 runs.

### 7 T data acquisition

High spatial resolution MRI data were collected on a 7T Philips Achieva MR system (Philips Healthcare, Best, Netherlands) using a volume transmit coil and a 32-channel receive coil (Nova Medical, Wilmington, MA), installed at Sir Peter Mansfield Imaging Centre, University of Nottingham, UK. To minimize head motion, we stabilized participants with a customized MR-compatible vacuum pillow (B.u.W. Schmidt) and foam padding.

Functional MRI data were obtained using T2*-weighted, gradient echo, three-dimensional echo-planar imaging with the following parameters: TR = 88 ms, TE = 28 ms, FA = 22°, EPI factor 45. We used parallel imaging (SENSE reduction factor of 2.35 in the anterior-posterior (AP) direction and 2 in the foot-head (FH) direction)^44^. Each complete volume acquisition (dynamic scan time) took 4 s. The spatial resolution of the acquired data was 0.75 mm isotropic and the field of view (FOV) of the imaging volume was 74 mm, 36 mm, and 154 mm in the anterior-posterior, foot-head, and right-left directions respectively. The reduced FOV in the phase encoding direction (AP) required the use of outer-volume suppression to prevent signal fold-over. Simulations to estimate the point-spread function for the imaging parameters used were performed (Matlab 8.0, Mathworks, Natick, MA) assuming a T2* of 25 ms for gray matter, and resulted in a full-width-half-height of less than 1.4 voxels, giving an effective resolution of approximately 1 mm in the phase encoding direction.

An image-based shimming approach was used^45-47^; this required the acquisition of a B_0_ map prior to the fMRI acquisition, and 2^nd^ order shimming was restricted to a cuboidal region covering the calcarine sulcus. These shim settings were then fixed for the entire functional MRI experiment to prevent scan-to-scan changes in local geometric image distortions.

To simplify alignment of the functional MRI data to the whole head anatomy (MPRAGE), we also obtained T_1_-weighted anatomical images for each subject at the end of their scan session. These images had the same slice prescription and coverage as the functional data. MPRAGE sequence parameters were: linear phase encoding order, TE = 2.1 ms, TR = 14 ms, FA = 10°, TI = 960 ms, 2 averages, SENSE acceleration factor 2.5, ~2 min acquisition time). In addition, we obtained high-resolution, T2*-weighted axial images using 2D FLASH (0.25 × 0.25 × 1.5 mm^3^ resolution; TE = 20 ms, TR = 320 ms, FA = 32°).

### Phase encoded data analysis

Preliminary analysis of functional data was performed with a combination of software (mrTools, NYU, http://www.cns.nyu.edu/heegerlab/wiki/; VISTA, Stanford) running in Matlab 8.0 (Mathworks, Natick, MA) and FSL (FMRIB Software Library, FMRIB, Oxford, UK). Further analysis of the data performed with the R statistical software package and various toolboxes for this environment as detailed in the section “variogram analysis”.

Flattened representations of the occipital cortex were generated in order to locate the early visual cortical areas for each subject from MPRAGE images obtained at 3 T. Segmentation was performed anatomically using FreeSurfer (http://surfer.nmr.mgh.harvard.edu/). Gray matter smooth 3D surfaces were flattened at the occipital pole (radius 70, manifold distance) using mrTools and custom-written software (mrTools, NYU, http://www.cns.nyu.edu/heegerlab/wiki/). From these flattened cortical representations, retinotopic maps of the left and right visual cortex were made for all subjects. A correlation analysis was performed on the average of the 6 retinotopic mapping runs; this analysis calculates how well each voxels2019 time series correlates with the experimental design and therefore can be used to match visual areas to the regions of the visual field that they represent. Regions of interest corresponding to areas Vl, VZ, V3, and V3A were defined for all subjects.

Data from Experiment 2 (disparity waveforms) were analyzed using standard Fourier-based methods as for retinotopic mapping (mrTools) and displayed on flattened cortical representations. An initial detection of clusters was performed using a correlation analysis on an average of the 8 runs, which was then imported onto the flat maps to assess the clustering and distribution of activity across early visual areas for each subject. For this clustering measure, a he threshold measure involved producing a new image in which the value of each individual voxel represented the standard deviation of the surrounding 342 voxels (approximately 5 mm radius). Since the values were a measure of response phase, a cyclical metric, circular standard deviation was obtained with the following equation

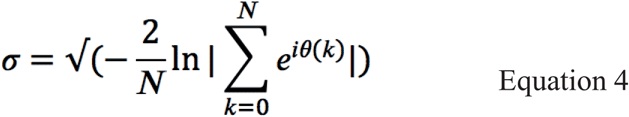

where θ is the phase of each voxel.

Thresholding this metric at 2 standard deviations ensures that only voxels with a similar phase are considered, essentially a type of smoothing. However, this method ensures individual voxels maintain their original value, which is necessary to investigate phase changes across the cortical thickness.

### Variogram analysis

In the cases we examine here, the imaging data is inherently discontinuous because of the thresholding effect of the stimulus coherence threshold. It is therefore more accurate to consider the thresholded data as a list of observations *Z*_*i*_ at points *(x*_*i*_*, Y*_*i*_*, z*_*i*_). This defines a collection of points to which the methodology of spatial point processes can be applied. It is possible to define first and second moments of the statistics, corresponding to local mean and local variance. To describe the spatial dependence of the measured quantities, spatial statistics defines a fundamental statistic called the variogram, which is the variance of the difference in measured values taken at two different locations within the sampling field. We begin initially with the calculation of the variogram on unfolded ‘flat maps’ of the cortical sheet so that the representation of the data is two-dimensional, *Z*_*i*_ = *(x*_*i*_*, y*_*i*_).Thus for two observations *Z_i_* and *Z_j_* at points *u* = *(x*_*j*_*, y*_*j*_) and *v* = *(x*_*j*_*, y*_*j*_) then the variogram is given by

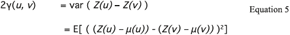

where E [] is the expected value of the quantity and *μ* is the mean value and *γ* is the semi-variogram. In the case where the mean value *μ* is constant across the imaging field, this reduces to

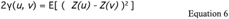

Further, where the underlying data values are stationary, the variogram depends only on the separation between *u* and *v,* say *w.*

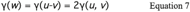

Modeling of noise patterns was carried out using the R statistical software system [R **version 3. 1. 2** (2014-10-31)]^48^ with several extension packages (notably ‘`fields`’ **version 8. 2-1** and ‘`gs tat`’ **version 1.0-21** available from the CRAN source http://cran.r-project.org). Analysis of the empirical distribution of variance in the MRI images was performed using the vgram function from the fields package.

The empirical variogram was fit with a cumulative Gaussian model with three parameters (variance at large values of *w* in Equation 3, so-called “sill” variance; variance for *w =* 0, so-called “nugget” variance; and the range parameter, specified by the steepness of the cumulative Gaussian). Of these parameters, the most interesting is the range, as this specifies the spatial distance over which the samples of fMRI signal are correlated with the values at the reference point.

Values of the range parameter were extracted for every experimental condition from each subject for each of their identified visual cortical areas. These calculations of the range were calculated separately for pixels in the flat-map with high coherence (> 0.3) with the changing disparity waveform and for pixels with low coherence (<= 0.3). The low coherence range values form an estimate of the baseline level of spatial correlations within the fMRI signals across the cortical surface. These correlations limit the spatial resolving power of the MRI signal^8^ and the spatial resolution estimated from the values of the range parameter at low coherence match closely independent estimates of the spatial resolution of fMRI signals at 7T field strength^49^. As the cortical activations are specified by a complex quantity with amplitude and phase, the activities were first converted to sine and cosine components before the variogram analysis and curve-fitting were performed (see Supplementary Figure 2) before the noise synthesis was performed.

Returning to the question of whether to assess the variogram in two or three spatial dimensions, we first note that the organization of functional columns across the cortical surface is inherently two-dimensional. For this reason above all, our focus for this investigation was on calculations of the variogram across the two-dimensional cortical sheet. Nonetheless, it is also possible to calculate the three-dimensional variogram for data represented in native imaging space. This has one advantage over two-dimensional calculations on flat maps, in that the process of unfolding to form the flat map has the potential to introduce local spatial distortions. These distortions alter the distance measurement between neighboring pixels in the flat-map and hence the variogram calculation. It is hard to see that this will affect the comparison of functional data at high and low stimulus coherences, since any distortions are identical in both data sets because they are both specified on the same flat-map. Nonetheless there is the possibility that absolute numerical values of the range parameter for the variogram could be affected by the distortions inserted by the unfolding process.

Accordingly, we went ahead and made calculations of the 3-D variogram structure, including the fitting of a range parameter. There are some limitations in carrying out the calculation of the variogram in native imaging space. The most significant of these are the inclusion of white-matter voxels in the calculated activity profiles and errors introduced by the close apposition of grey-matter voxels from unrelated areas of cortex that may occur within cortical sulci. These limitations can be potentially addressed more fully with more detailed analysis, but for now we restrict comparisons to the extrastriate cortical areas V2, V3 and V3A, thereby avoiding any confounds from V1 due to the calcarine sulcus. Consistent with the view that local spatial distortions may affect the absolute values of range parameter, the average value for the range parameter for the 3-D variogram of these extrastriate cortical areas is smaller (ratio of 2-D range to 3-D range = 1.31; low stimulus coherence condition for both cases).

These values are calculated on the assumption of isotropy in 3-D native imaging space and the 2-D flat map. We note that there is considerable interest in the question of functional imaging measurements across the cortical layers from pial surface to white matter. As the orientation of the cortical grey matter varies considerably relative to the coordinate frame for native imaging space, the presumption of isotropy is unlikely to be fully correct, even if the data analysis is stringently restricted only to voxels that encompass grey matter.

### Disparity topography analysis

For each spatial array, whether phases of simulated noise or high coherence data, the same procedure was followed. A set of four cardinal directions on the data field was chosen, being 0, 45, 90, and 135 degrees. Analysis proceeded along all available lines in the 2D data field that lay parallel to a cardinal direction. Along each line, the phases were unwrapped to ensure no artificial breaks in the phase distribution and then differentiated. The differentiated phase lines were assessed using a simple run length analysis, which reported the incidence of a consistent positive or negative sequence of values and the length of the run. Each identified run-length was accumulated to form a frequency distribution of run-lengths. This was done separately for the simulated phases from the noise distribution and the high-coherence data. These two frequency distributions were compared using a Kolmogorov-Smirnov test (two-sample test with correction for discrete levels, using `dgof` package in R) to determine whether the true data at high coherences could have been generated by chance sampling from the distribution for noise distributions that matched the spatial correlation structure of the low-coherence data. All these Kolmogorov-Smirnov tests were highly significant (p << 0.001), indicating that the high coherence fMRI data carries spatial characteristics that are absent from the low coherence fMRI signals.

### Mark correlation analysis of disparity topography

In the language of spatial statistics, the data here consist of a ‘mark’ (i.e. a value of a property, in our case phase angle) at a location *x,y.* The mark correlation function (Equation 2) compares the expected value *E* of the actual distribution of marks in the numerator against the null hypothesis of a random association between marks and spatial location. A suitable function for comparing the difference of two phases is the absolute difference (Equation 3): a number of alternative comparison functions were explored with no substantial difference to the results. The expected value of the correlation function is 1, under the assumption of no link between the distance between two points and the difference of the mark values (difference of phases). Values less than 1 imply a positive association (phase differences are smaller than average if points are closer) and those greater than 1 imply a negative association (phase differences are greater than average if points are closer). The mark correlograms in Figs 4 were calculated according to equations 2 and 3, using the markcorr function from spatstat (version 1.48.0) package^39^ in the R environment.

## Acknowledgements

This work was supported by a grant to HB & AJP from the Medical Research Council (G0802171) and a Royal Society University Research Fellowship to HB.

## Supplementary Figures

**Supplementary Figure 1:**
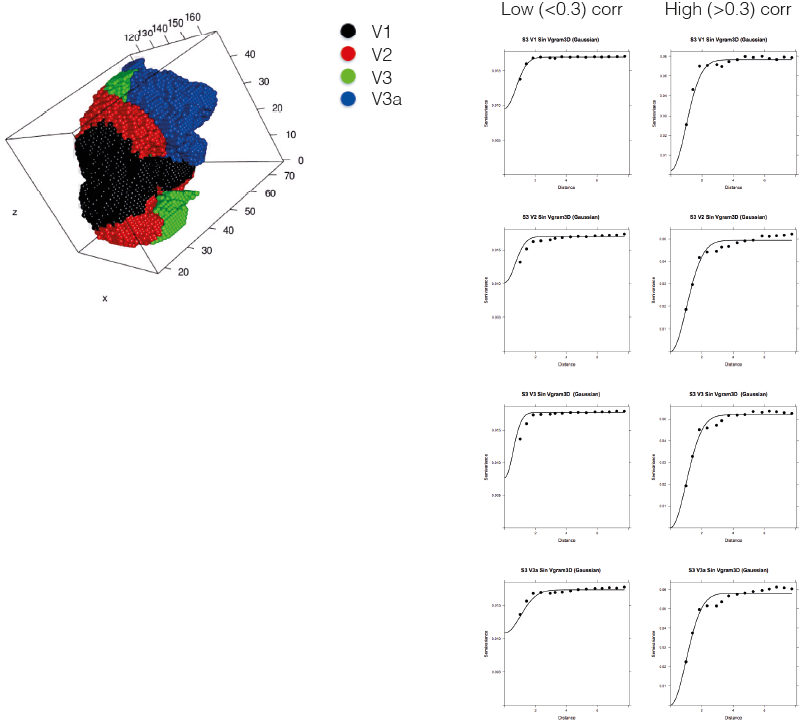
calculation of variogram of experimental data in native imaging space under the assumption of radial, spherical symmetry. Three-dimensional rendering of locations of data in native imaging space from 4 cortical areas in occipital cortex in upper left panel: V1 (black), V2 (red), V3 (green) and V3A (blue). Right hand panels show calculation of variogram in 3D space for sine component of amplitude/phase data (see Supplementary Figure 2) of subject S3. Low coherence data show same pattern of a rise in the variogram as a function of distance until a stable value is reached for longer-range data samples. High coherence data show the same deviation from the simple Gaussian model, implying more correlation structure at greater separations of sampling points. An ANOVA with low vs high as a main effect gives F(1,126) = 17.025; P < 6.654e-05. The mean value of range averaged across all three subjects with repeated data was very similar for the low coherence voxels for all four cortical areas (V1: 1.29; V2: 1.24; V3: 1.26; V3A: 1.27; Grand Mean: 1.265; units of sampling distance, equivalent to 0.95mm half-width at half-height for a Gaussian sphere).

**Supplementary Figure 2:**
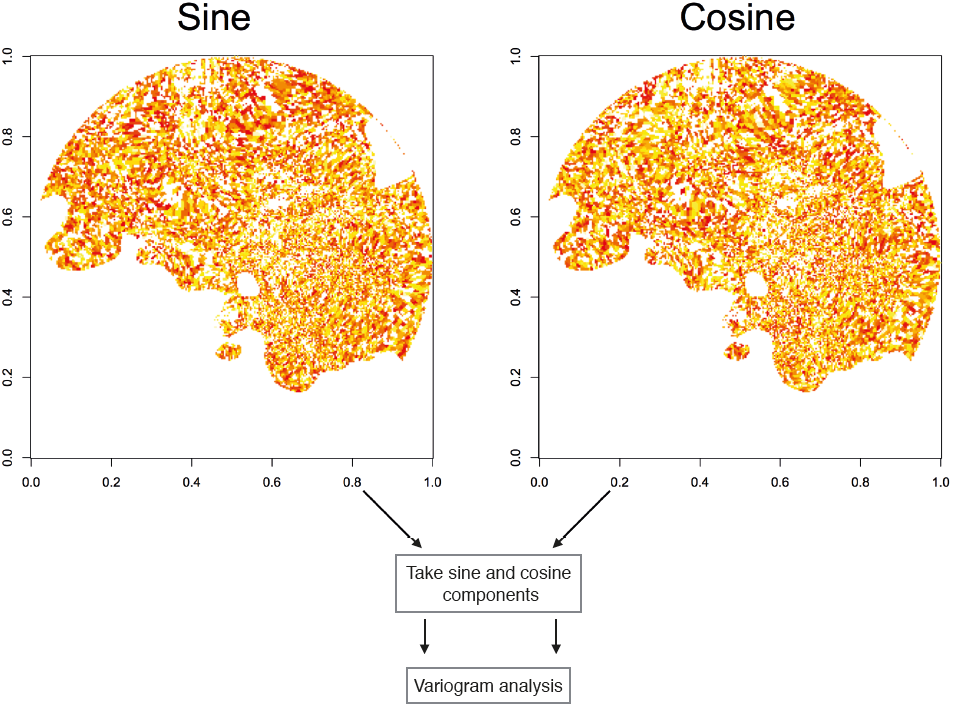
Low coherence signal for subject S3 represented as sine and cosine components, prior to variogram analysis

**Supplementary Figure 3:**
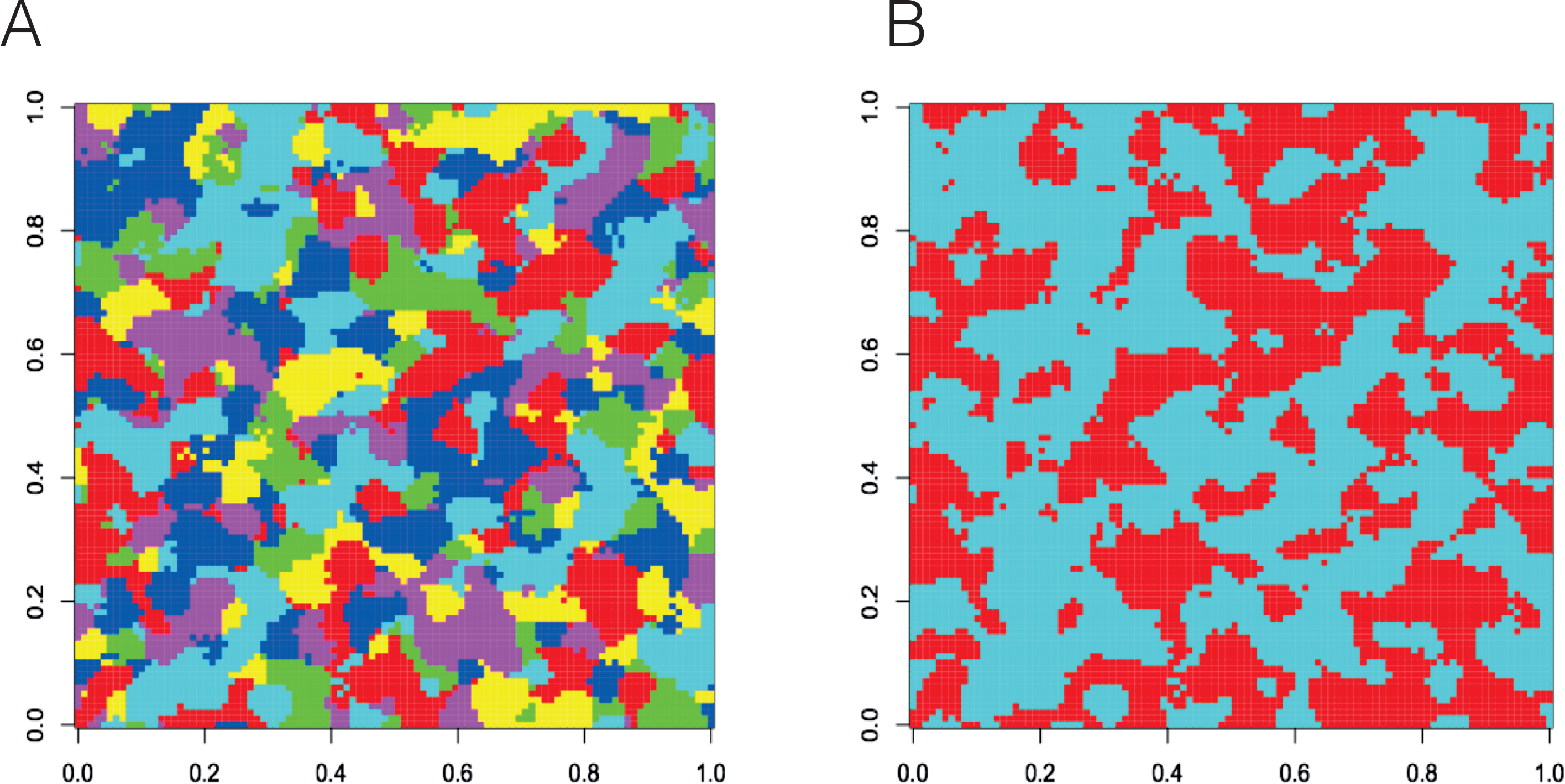
pattern of apparent stimulus-specific zones of cortical map induced by spatial correlations of signal across the cortical surface. A: six independent samples of spatially correlated noise were generated and each zone of the map in A was color coded according to which of the six samples had the greatest response at that pixel. B: similar coding of spatially correlated noise with two samples rather than six. The spatial correlations within the noise samples result in a local dominance of the synthetic cortical map. The size and shape of these dominant regions depend on the correlation distance and the number of categories that are assigned (2 vs 6 in this case). Although these patterns are generated entirely by noise, they have some similarity with patterns of cortical columns reported in the imaging literature. Note that each new set of noise samples will result in a pattern with a different detailed distribution of dominant regions, although the size and shape of the dominant regions will be similar for each generated pattern.

**Supplementary Figure 4:**
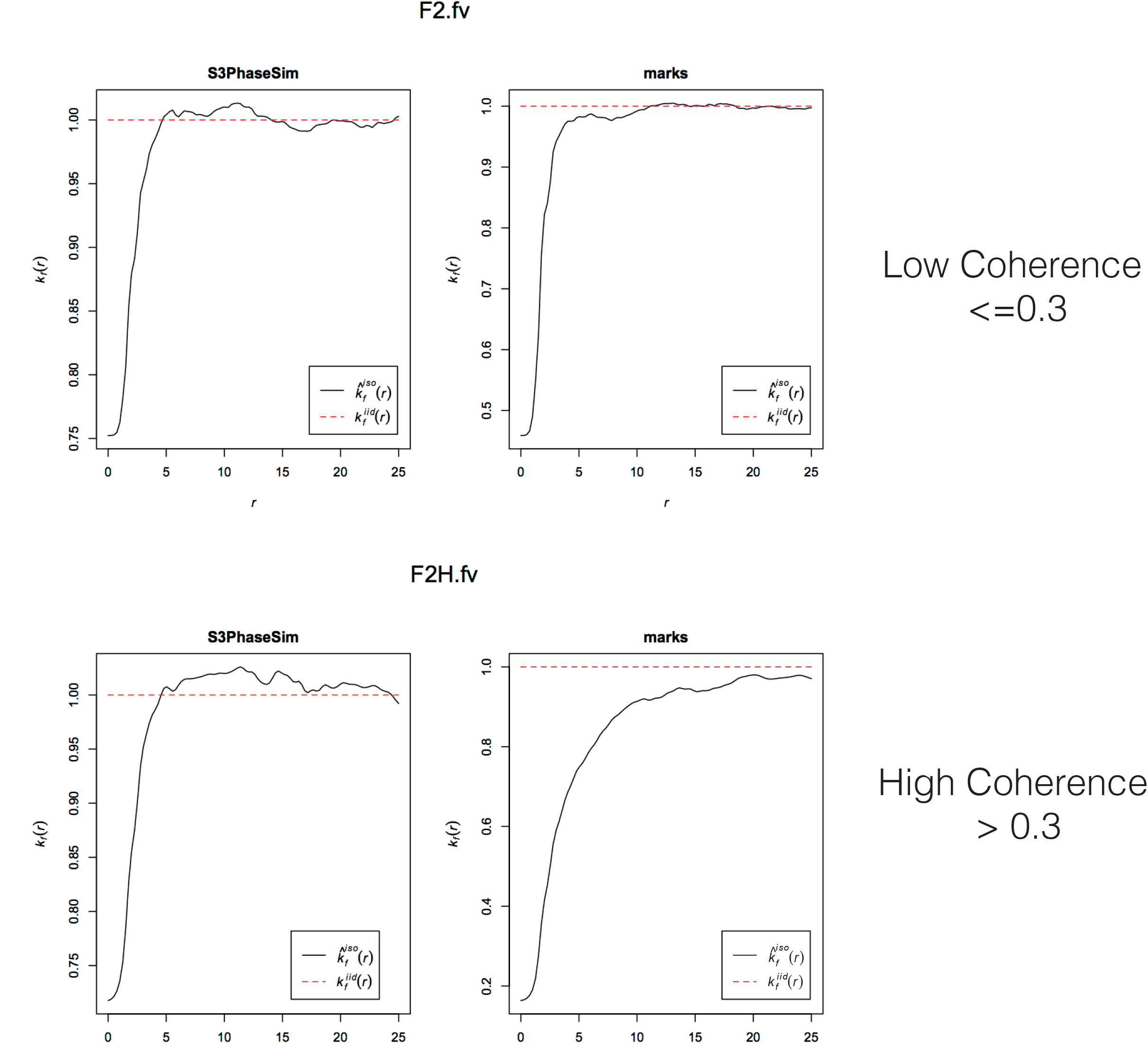
Comparing the mark correlation function for values of phase for simulation (left panels, “PhaseSim”) and experimental data (right panels, “marks”). Each panel plots the value of mark correlation (y axis) against the cortical distance (in units of sampling distance — 0.75mm separation): see Equation 2 main paper for specification of mark correlation calculation. A value of 1 in the mark correlation means no correlation, whilst values less than 1 imply neighboring samples are more similar than chance association. The upper panel shows low coherence data, while the lower panel shows high coherence data. Simulations adequately capture the spatial structure of the phase distributions of low coherence data, which forms a noisy background against which the high coherence data must be detected. The high coherence experimental data (bottom right) show a positive association of phase values between neighboring sample points out to some 15-16 units, equivalent to interactions over a distance of 11-12mm across the cortical map.

**Supplementary Figure 5:**
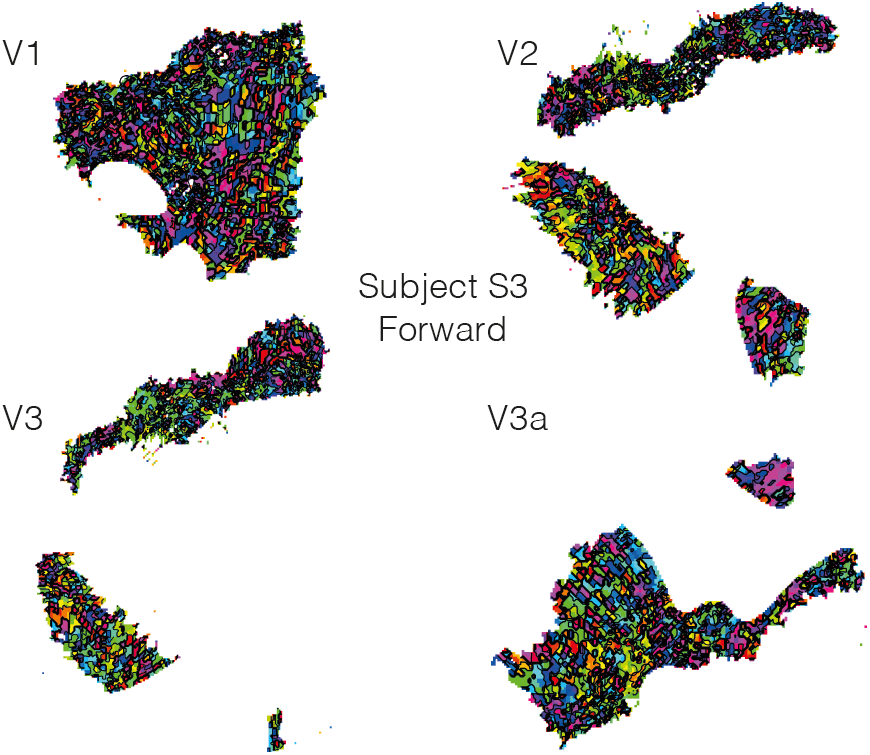
Color-coded contour maps of preferred disparity phase for all (high and low coherence) pixels on flat maps of cortical areas V1, V2, V3, and V3A for subject S3 Forward condition.

**Supplementary Figure 6:**
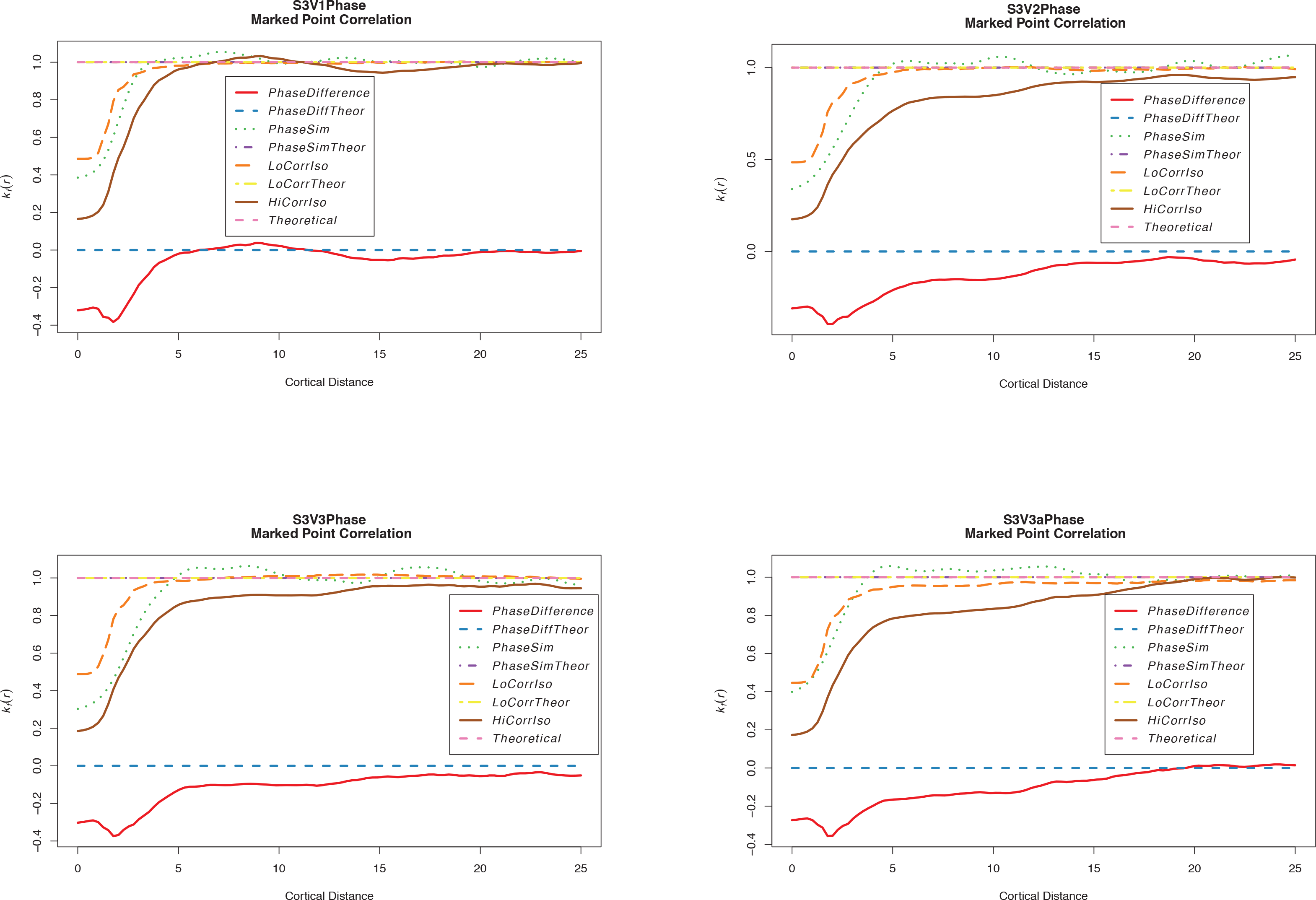
Marked point correlation plots for the four cortical areas shown in Supplementary Figure 5 for subject S3 Forward condition

**Supplementary Figure 7:**
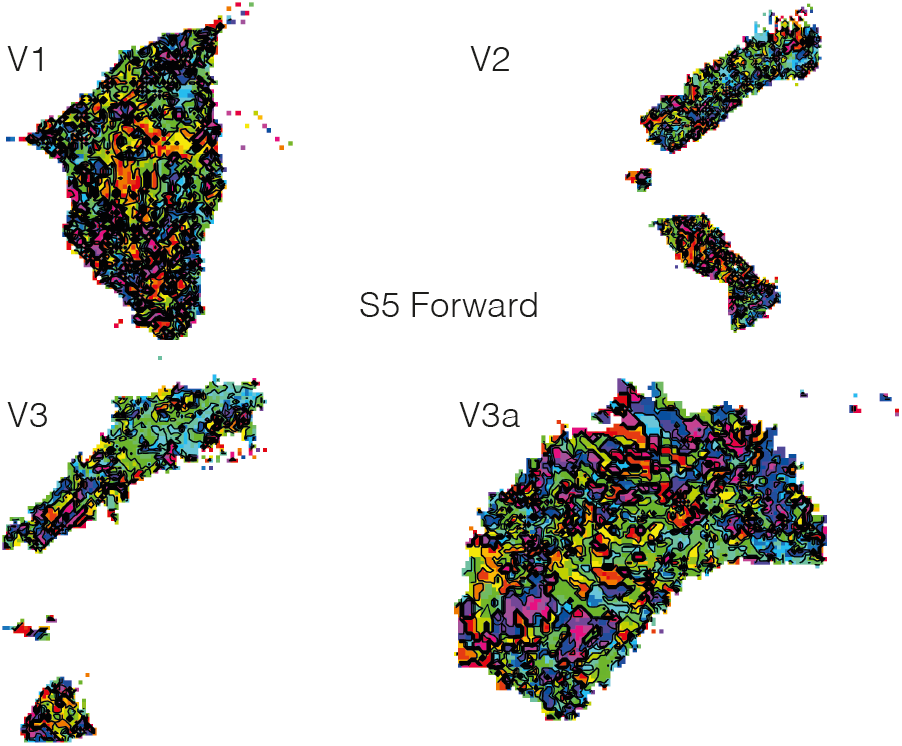
Color-coded contour maps of preferred disparity phase for all (high and low coherence) pixels on flat maps of cortical areas V1, V2, V3, and V3A for subject S5 Forward condition.

**Supplementary Figure 8:**
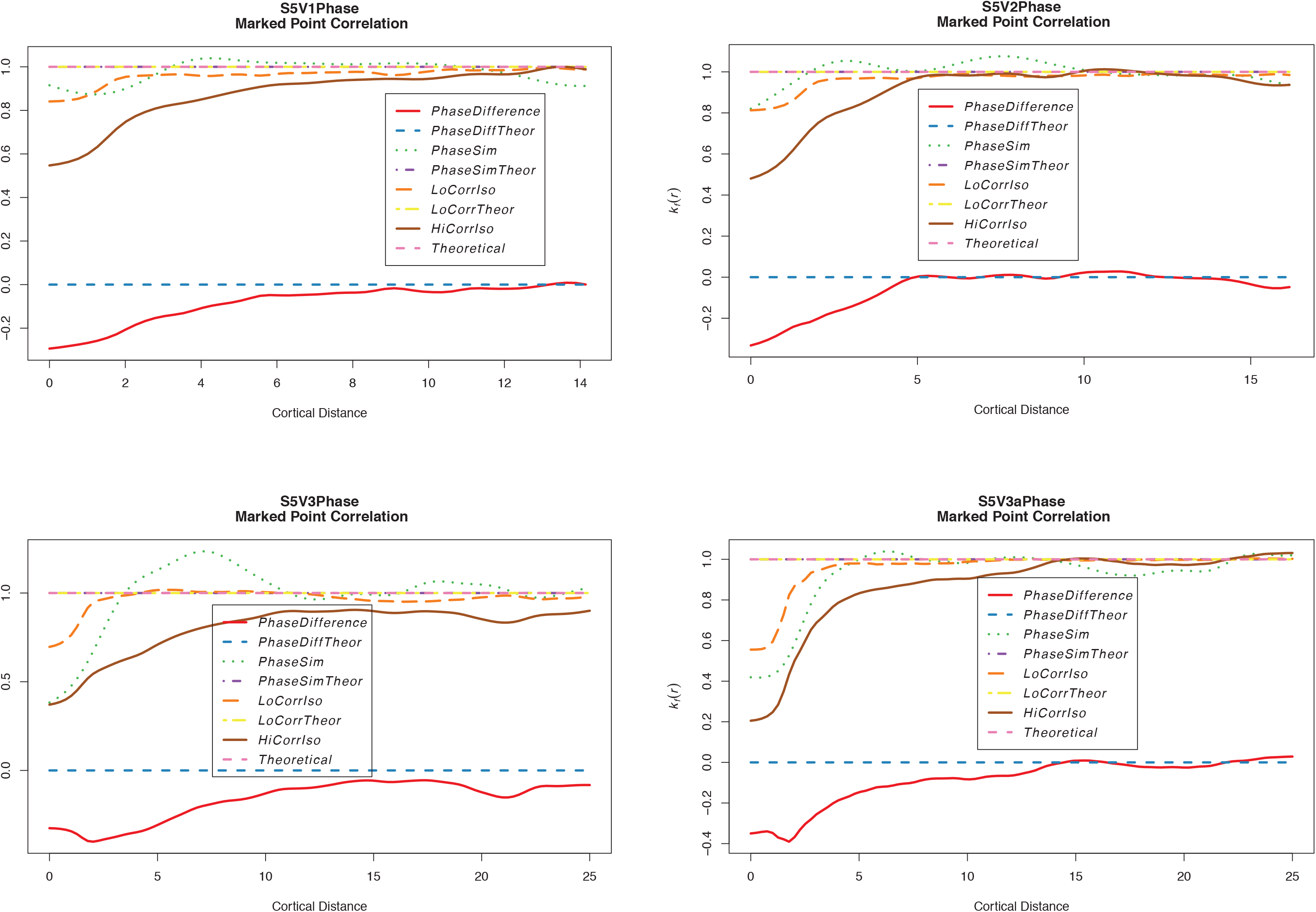
Marked point correlation plots for the four cortical areas shown in Supplementary Figure 7 for subject S5 Forward condition

**Supplementary Figure 9:**
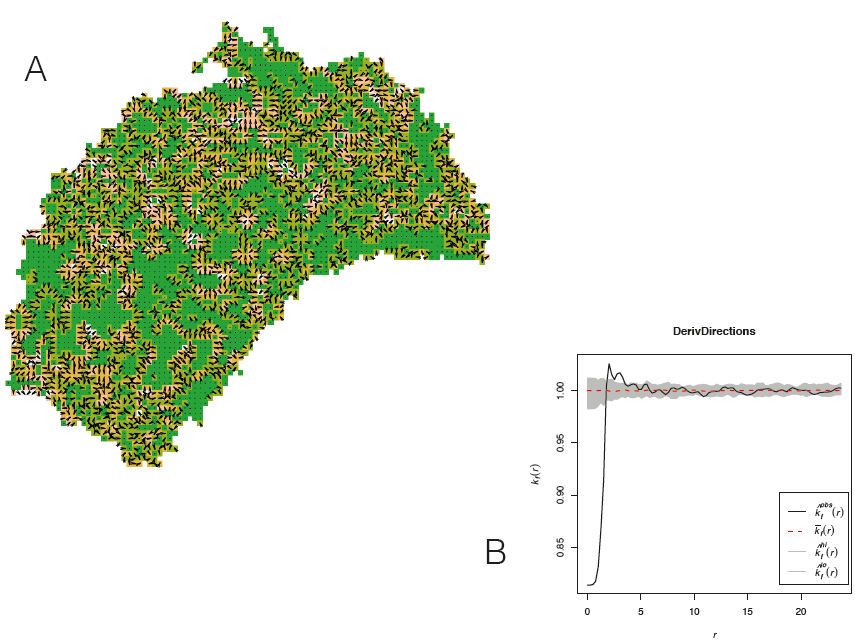
Testing for systematic changes in preferred disparity across a cortical region. A: shows the amplitude of the two-dimensional derivative of the disparity field, color-coded as in Figures 4 and 5 (main paper). Data from subject S5; Reverse condition. Superimposed arrows show direction of maximum gradient of the differential disparity map, thus showing the direction of greatest change of disparity at a particular point on the map. B: The local directions were subjected to a mark correlation analysis to reveal whether nearby points have similar or different changes of disparity preference. Any local similarity is short range, with a modest tendency to have different directions of slope at about 3 cortical units distance (0.75mm voxel steps). The grey shading shows a bootstrap estimate of the point-wise 95% range of expected data under the null hypothesis of no relationship between cortical distance and value of the mark (in this case direction of disparity change).

